# Wnt signaling determines body axis polarity in regenerating *Hydra* tissue

**DOI:** 10.1101/2020.01.22.915702

**Authors:** Rui Wang, Robert E. Steele, Eva-Maria S. Collins

**Author notes:** **Corresponding Authors**: Eva-Maria S. Collins.

## Abstract

How an animal establishes its body axis is a fundamental question in developmental biology. The freshwater cnidarian *Hydra* is an attractive model for studying axis formation because it is radially symmetric, with a single oral-aboral axis. It was recently proposed that the orientation of the new body axis in a regenerating *Hydra* is determined by the oral-aboral orientation of the actin-myosin contractile processes (myonemes) in the parent animal’s outer epithelial layer. However, because the myonemes are not known to possess polarity, it remained unclear how the oral-aboral polarity of the axis would be defined. As Wnt signaling is known to maintain axis polarity in *Hydra* and bilaterians, we hypothesized that it plays a role in axis specification in excised *Hydra* tissue pieces. We tested this hypothesis using pharmacological perturbations and novel grafting experiments to set Wnt-derived signals and myoneme orientation perpendicular to each other to determine which controls axis formation. Our results demonstrate that Wnt signaling is the dominant encoder of axis information, in line with its highly conserved role in anterior-posterior patterning.

## Main text

Coordinated interplay of chemical and mechanical signaling is critical for morphogenesis and patterning in metazoans (e.g. (1)). However, their relative contributions are context-dependent and can be difficult to determine in a living animal. The freshwater cnidarian *Hydra* has long been popular for studying patterning (2) and is now being developed as a model for biomechanical studies (3–6), as it enables *in vivo* examination of the mechanochemical basis of pattern formation.

*Hydra* can regenerate from small pieces of tissue excised from the body column and it has long been thought that sufficiently small pieces lose parental axis information and undergo *de novo* axis specification (7). However, a recent study (3) asserts that excised *Hydra* tissue pieces inherit the parental body axis through actin-myosin contractile elements (myonemes) oriented parallel to the body axis in the ectoderm. While this work (3) shows a strong correlation between myoneme orientation and axis orientation, it does not establish a causal link. Here, we use grafting experiments and manipulation of canonical Wnt signaling via alsterpaullone (ALP) treatment (Methods) to demonstrate that Wnt signaling defines body axis polarity and directs myoneme orientation in regenerating *Hydra* tissue pieces.

To test oral-aboral polarity inheritance, we generated oral-aboral grafts using animals of two differently labeled *Hydra* strains, a fluorescently labeled strain and an unlabeled strain, bisected perpendicular to the body axis (8). Excising tissue pieces from the subsequent graft interface allowed us to determine the orientation of the regenerated axis relative to the axis of the grafted animal (**Fig.1A**). Axis polarity was assayed by determining whether the strain giving rise to the regenerated head was the same as that in the oral half of the grafted animal (**Fig.1Bi, ii**). Of these grafts, 37/40 retained the polarity of the grafted animal and 3/40 reversed it (**Fig.1C**). We therefore conclude that axis and polarity information are inherited in small tissue pieces, in agreement with (3). Next, we perturbed Wnt signaling by grafting the lower halves of ALP-treated polyps to upper halves of untreated animals. *Hydra* expresses Wnt in the head organizer at the tip of the hypostome, while ALP causes ectopic Wnt signaling throughout the animal (9). Grafted animals thus have normal Wnt signaling in the oral half, and ectopic Wnt signaling in the aboral half (**Fig.1Biii**). Of these ALP+ grafts, 6/50 retained the original polarity while 44/50 reversed it, a significant difference from untreated (ALP-) grafts (p=1.896e-15) (**Fig.1C**). Myoneme organization at the graft interface was visualized using phalloidin (5) and appeared normal, with ectodermal myonemes parallel to the body axis in both the presence and absence of alsterpaullone treatment (**Fig.1D**). Together, these data demonstrate that while tissue pieces retain their original myoneme organization, their polarity can be reversed by ectopic Wnt signaling in the lower half of the grafted animal.

**Figure 1.**
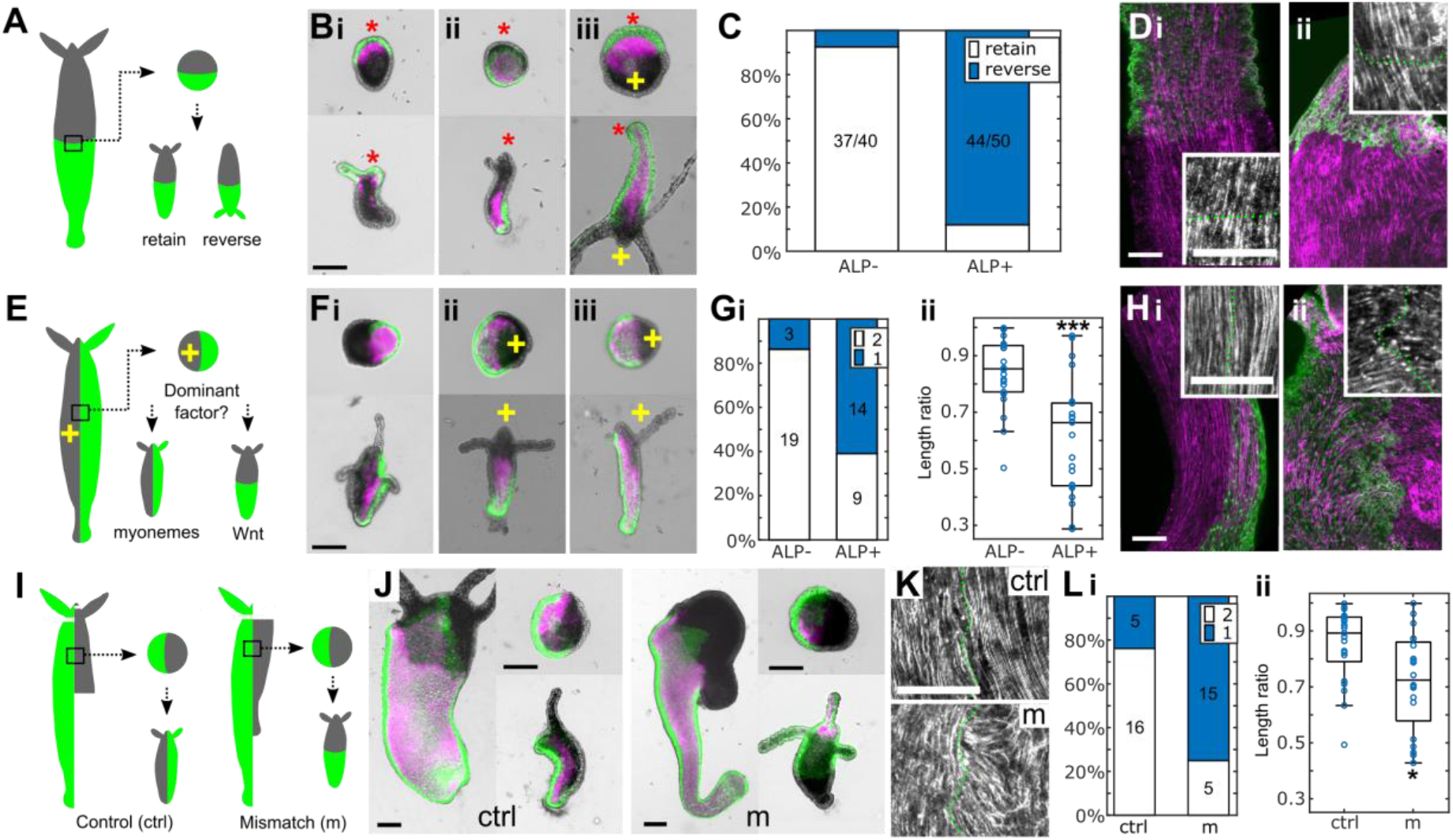
Wnt signaling encodes axis polarity. A. Experimental schematic of oral/aboral grafting, showing grafted animal, cut tissue piece, and observed outcomes. B. Three examples of tissue pieces from the interface of oral/aboral grafts, before (top) and after (bottom) regeneration. Red asterisks indicate direction of parental head where known. Yellow crosses indicate ALP treated tissue. This notation is also used in E and F. (i) Control with fluorescent watermelon (WM) head, (ii) control with WM foot and (iii) animal with WM head and ALP-treated foot (ALP+) and WM head. C. Axis polarity results for control (ALP-) and ALP+ oral/aboral grafts (p=1.896e-15). D. Representative phalloidin stains of oral/aboral grafts, insets show graft interface at higher magnification. Scale bars 100 μm. i. ALP-, ii. ALP+. E. Experimental schematic of lateral graft, showing grafted animal, cut tissue piece, and possible outcomes. F. Examples of tissue pieces from the interface of lateral grafts before (top) and after (bottom) regeneration, showing range of observed outcomes. i. ALP- control. ii, iii. ALP+. G. Control vs. ALP+ lateral grafts. i. Number of strains in head (p= 0.0018). ii. Interface/body length ratio (p= 1.5651e-04). *** indicates p<0.001. H. Phalloidin stains of lateral grafts. i. ALP-, ii. ALP+. H. ALP- vs. ALP+ lateral grafts. I. Schematics of control (ctrl) and mismatch (m) quadrant lateral grafts. J. Representative images of control and mismatch quadrant lateral grafts. Scale bars 200 μm. K. Phalloidin stains of quadrant graft interfaces. L. Control vs. mismatch quadrant grafts. i. Number of strains in head (p= 0.0017). ii. Interface/body length ratio (p= 0.011). * indicates p<0.05.

Because ectodermal myonemes and Wnt signaling are both oriented parallel to the body axis, our results from the oral-aboral grafts cannot distinguish between their contributions to the inheritance of polarity and oral-aboral axis orientation. To address this question, it was necessary to set myoneme orientation and Wnt signaling in direct conflict with each other. We engineered such an outcome using lateral grafts (**Fig.1E, F**) wherein one half has been treated with ALP and one is untreated. This creates an ALP+ chimera in which Wnt signaling is strongest in the ALP-treated half of the animal, perpendicular to the body axis. Tissue pieces cut from the interface of ALP+ chimeras (n=23) were compared to those originating from lateral grafts of untreated animals (n=22). If initial myoneme orientation determined axis orientation, as proposed in (3), tissue pieces from the interface of an ALP+ lateral graft should regenerate with a split between labeled and unlabeled tissue parallel to the body axis as seen in control lateral grafts. Alternatively, if Wnt signaling determines axis orientation, they should show ALP+ tissue in the head. Two scoring metrics were used to account for experimental variability caused by the grafting procedure: number of tissue types (ALP treated/untreated) in the head and the ratio of the length of the interface between the two tissue types to overall body length (Methods). We found statistically significant differences between ALP+ and ALP- lateral grafts (p<0.05; **Fig.1G**), demonstrating that Wnt signaling defines oral-aboral polarity and overrides the role of myonemes in setting body axis orientation. Moreover, while alignment of the ectodermal myonemes was parallel to the body axis in untreated control grafts, we observed a degree of myoneme reorientation perpendicular to the body axis in ALP+ grafts at < 24 hrs post-grafting (**Fig.1H**), suggesting that Wnt signaling can redirect myoneme orientation.

One could argue that under physiologically relevant conditions, the body axis is set by myoneme orientation, and that the unnaturally high level of Wnt signaling induced by ALP treatment overrides mechanical signals. However, previous experiments suggest that myoneme orientation is a consequence of Wnt signaling, even under physiological conditions. For example, during budding *Hydra* Wnts are consistently expressed as early as stage 1 (9), when remodeling of myonemes has not yet occurred (11). Furthermore a head organizer grafted into the body column induces an ectopic body axis (see e.g.(10)), which requires the remodeling of surrounding myonemes similar to what is observed during budding. This is consistent with our observation that myonemes reorient perpendicular to the graft interface in ALP+ lateral grafts in less than 24h (**Fig.1Gii**).

To directly test whether physiological Wnt signaling levels can override preexisting myoneme orientation, we carried out “quadrant” lateral grafts (**Fig.1I,J**). These were created either by matching the donor quadrant to the recipient half (control), or by using a mismatch to appose tissue from the oral and aboral ends of two animals. Control grafts with no mismatch show normal myoneme alignment, while mismatch grafts show signs of myoneme reorientation, as observed in ALP+ lateral grafts (**Fig.1K**). Tissue pieces cut from the interface of quadrant grafts without a mismatch were significantly different from those with a mismatch in both scoring criteria (**Fig.1L**). In addition, mismatch quadrant grafts were similar to ALP+ lateral grafts (strains in head, ratio = 0.4566, 0.7065) while control quadrant grafts were similar to ALP- lateral grafts (0.3528, 0.0735). This indicates that quadrant grafting does not inherently alter regeneration behavior, but physiologically relevant differences in Wnt signaling can override structural cues in establishing a new axis. Thus, our work explicitly shows that Wnt signaling encodes both the orientation and the polarity of the oral-aboral axis in *Hydra*.

## Materials and Methods

*Hydra* were maintained using standard methods (12). Oral/aboral grafts (as in (8)) and lateral grafts were performed using transgenic watermelon (WM) (13) and wild type AEP strains. The position (oral/aboral) of the lineages in the grafts was random, to account for possible strain differences. Animals were treated with alsterpaullone (ALP) for 2 days as in (9). Quadrant lateral grafts were made by fixing one half of an animal on needles, followed by adding a quarter of a second animal to the needle fixing the aboral end. The tip of the hypostome was excised from all pieces to remove existing head organizers and delay reestablishment of a normal gradient. Imaging using confocal microscopy was performed as described in (8). Images of oral/aboral grafts were manually scored for polarity retention by assessing if the lineage represented in the head in the regenerated animal matched that in the original animal. Lateral graft regeneration was scored by two metrics: the number of strains represented in the regenerated head above the tentacle ring, and the ratio of the length of the interface between strains to the length of the animal measured down the midline from hypostome to foot. All measurements were conducted using Fiji (14). For all grafts, only regenerated animals able to feed after 4 days were included in the analysis. Raw data are available upon request. Statistical significance was determined using Fisher’s Exact Test for polarity retention and number of tissue types in the head, and the Mann-Whitney U Test for interface ratio. Statistical analysis using multiple metrics is necessary as lateral grafts frequently showed uneven interfaces or slightly inconsistent results, caused by the intrinsic difficulty of this experimental procedure and the impossibility of matching the biochemical signaling of two animals without the ability to visualize such signaling directly.

## Acknowledgements

We thank S. Martin for help with *Hydra* care and discussions, S. Guan and T. Goel for help with experiments, and Drs. D. Ireland and K. Willert for comments on the manuscript. This work was supported by NSF grant CMMI-1463572, the Research Corporation for Science Advancement, and the Gordon and Betty Moore foundation.

## References

1. Mammoto, T., and D.E. Ingber. 2010. Mechanical control of tissue and organ development. Development. 137: 1407–20.

2. Meinhardt, H. 2012. Modeling pattern formation in hydra: a route to understanding essential steps in development. Int. J. Dev. Biol. 56: 447–462.

3. Livshits, A., L. Shani-Zerbib, Y. Maroudas-Sacks, E. Braun, and K. Keren. 2017. Structural Inheritance of the Actin Cytoskeletal Organization Determines the Body Axis in Regenerating Hydra. Cell Rep. 18: 1410–1421.

4. Mercker, M., A. Köthe, and A. Marciniak-Czochra. 2015. Mechanochemical symmetry breaking in Hydra aggregates. Biophys. J. 108: 2396–2407.

5. Wang, R., T. Goel, K. Khazoyan, Z. Sabry, H.J. Quan, P.H. Diamond, and E.-M.S. Collins. 2019. Mouth Function Determines the Shape Oscillation Pattern in Regenerating Hydra Tissue Spheres. Biophys. J. 117: 1145–1155.

6. Cochet-Escartin, O., T.T. Locke, W.H. Shi, R.E. Steele, and E.-M.S. Collins. 2017. Physical Mechanisms Driving Cell Sorting in Hydra. Biophys. J. 113: 2827–2841.

7. Soriano, J., C. Colombo, and A. Ott. 2006. Hydra molecular network reaches criticality at the symmetry-breaking axis-defining moment. Phys. Rev. Lett. 97: 258102.

8. Goel, T., R. Wang, S. Martin, E. Lanphear, and E.M.S. Collins. 2019. Linalool acts as a fast and reversible anesthetic in Hydra. PLoS One. 14: 584946.

9. Broun, M., L. Gee, B. Reinhardt, and H.R. Bode. 2005. Formation of the head organizer in hydra involves the canonical Wnt pathway. Development. 132: 2907–16.

10. Browne, E.N. 1909. The production of new hydranths in Hydra by the insertion of small grafts. J. Exp. Zool. 7: 1–23.

11. Otto, J.J. 1977. Orientation and behavior of epithelial cell muscle processes during Hydra budding. J. Exp. Zool. 202: 307–321.

12. Lenhoff, H.M., and R.D. Brown. 1970. Mass culture of hydra: an improved method and its application to other aquatic invertebrates. Lab. Anim. 4: 139–154.

13. Juliano, C.E., H. Lin, and R.E. Steele. 2014. Generation of Transgenic Hydra by Embryo Microinjection. J. Vis. Exp. 91: e51888.

14. Schindelin, J., I. Arganda-Carreras, E. Frise, V. Kaynig, M. Longair, T. Pietzsch, S. Preibisch, C. Rueden, S. Saalfeld, B. Schmid, J.-Y. Tinevez, D.J. White, V. Hartenstein, K. Eliceiri, P. Tomancak, and A. Cardona. 2012. Fiji: an open-source platform for biological-image analysis. Nat. Methods. 9: 676–682.

